# BL-Hi-C reveals the 3D genome structure of *Brassica* crops with high efficiency and sensitivity

**DOI:** 10.1101/2023.07.19.549753

**Authors:** Lupeng Zhang, Ranze Zhao, Jianli Liang, Xu Cai, Lei Zhang, Huiling Guo, Zhicheng Zhang, Jian Wu, Xiaowu Wang

## Abstract

High-throughput chromatin conformation capture (Hi-C) technologies can be used to investigate the three-dimensional genomic structure of plants. However, complex protocol and high background noise in Hi-C hinder its practical application in plant 3D genomics. Here, we took the approach of modified Bridge Linker Hi-C technology (BL-Hi-C) to explore plant 3D landscape. We modified the BL-Hi-C method by simplifing nuclei extraction step. By using *Brassica rapa* and *Brassica oleracea*, BL-Hi-C showed higher signal value and lower background noise than Hi-C. The high sensitivity of BL-Hi-C was further demonstrated by its capacity to identify gene loops involving *BrFLC1*, *BrFLC2* and *BrFLC3* which were undetectable in Hi-C. BL-Hi-C also showed promising performance with input as low as 100 mg leaf tissue. By analyzing of the generated data from BL-Hi-C, we found that the simulated 3D genome structure of *B. rapa* leaf cells was Bouquet configuration. Our results showed that the modified BL-Hi-C is a powerful tool for the investigation of plants’ genomic organization, gene regulation, and genome assembly.

**IN A NUTSHELL:** *Background:* 3D genome structure play a critical role in regulating spatiotemporal gene expression. However, there is a lack of simple, efficient and sensitive Hi-C technique in plants.

*Question:* How to study plant 3D genomics more simple and efficientHow to detect plant chromatin loops more sensitive?

*Findings:* We showed that BL-Hi-C is more simple, efficient and sensitive than coventional Hi-C by using *Brassica rapa* and *Brassica oleracea*. Furthermore, BL-Hi-C demonstrated its high sensitivity by detecting gene loops involving *BrFLC1*, *BrFLC2*, and *BrFLC3* which could not be detected by Hi-C. In addition, BL-Hi-C demonstrated promising performance with inputs as low as 100 mg leaf tissue. By analyzing BL-Hi-C data, we found that the simulated 3D genome structure of *B. rapa* leaf cells was Bouquet configuration.

*Next steps:* How chromatin loops are formed and regulated gene expression are key questions to be answered in plants. Our dataset of BL-Hi-C will enable future investigations to improve our understanding of chromatin loops.

## INTRODUCTION

The organization of DNA into chromatin in the nucleus of eukaryotic cells affects transcription, DNA replication, and other nuclear functions (Sexton and Cavalli, 2015). Chromosome conformation capture (3C)-based methods have significantly advanced our understanding of the organization of chromosomes (Bonev and Cavalli, 2016). High-throughput chromosome conformation capture (Hi-C) can interrogate all contact loci simultaneously, resulting in an all-to-all genome-wide interaction map by high-throughput sequencing (Belton et al., 2012). The application of Hi-C in plant research provides valuable insights into 3D genome organization. Chromosome occupies a relatively limited nuclear region, designated a chromosome territory (CT) (Fransz and de Jong, 2011), which shows different morphologies such as Rabl, Rosette and Bouquet. Many plants exhibit compartments and domains similar to topologically associated domains (TADs) as observed in recent studies (Xie et al., 2019; Wang et al., 2021; Zhang et al., 2021; Liao et al., 2022). In *Arabidposis*, 3C and Hi-C experiment had already shown that the promoter of *FLC* makes contact with downstream of *FLC* (Crevillen et al., 2013; Liu et al., 2016). Recent studies showed that dynamic 3D chromatin architecture correlated with genetic variance among parents and contributed to heterosis in *B. napus* (Hu et al., 2022). In *Brassica*, genes with higher numbers of conserved noncoding sequences (CNSs) are more likely to contact distant genes (Zhang et al., 2023). Pan-3D genome analysis reveals the relationship between structural variation and functional differentiation in soybean and links chromatin structures to cotton fiber length (Wang et al., 2022a; Ni et al., 2023).

Although Hi-C technology holds great potential for uncovering the landscape of 3D genomes, the complex protocol and high background noise limit its broad application. To address these issues, several advanced Hi-C methods have recently been developed. Capture Hi-C uses specific probes to capture the fragments related to the target region (Jager et al., 2015). In HiChIP, the target protein-specific antibody precipitates the DNA–protein complex after digestion and ligation (Mumbach et al., 2016). Digestion-ligation-only Hi-C (DLO-Hi-C) uses two rounds of digestion and ligation to complete the main experimental procedure (Lin et al., 2018). Hi-C on accessible regulatory DNA (HiCAR) (Wei et al., 2022) and ChIATAC (Chai et al., 2023) utilize Tn5 transposase and chromatin proximity ligation to analyze open-chromatin-anchored interactions. The application of these advanced Hi-C techniques in mammals has led to the discovery of cis-regulatory elements that regulate gene expression through remote interactions, thus extending the understanding of 3D genomics (Martin et al., 2015; Giambartolomei et al., 2021; Wei et al., 2022; Chai et al., 2023).

Bridge Linker-Hi-C (BL-Hi-C) technology has emerged as a powerful tool for exploring the chromatin architecture of the genome with a more straightforward procedure and reduced noise than other Hi-C technologies (Liang et al., 2017). This is due to the linker ligation strategy used in BL-Hi-C, which enables the digested chromatin to be ligated with the linker. This strategy is also employed in ChIATAC and CAP-C (You et al., 2021; Chai et al., 2023). In mammalian cells, BL-Hi-C has been instrumental in identifying key transcription factors that play a critical role in 3D genome organization, such as myogenic differentiation 1 (MyoD) (Wang et al., 2022b), and CCCTC-binding factor (CTCF) (Song et al., 2022). While BL-Hi-C has shown great potential in detecting 3D genome structure, its application in plant research is yet to be reported (Pei et al., 2021).

Traditional Hi-C is the main method for investigating the 3D genome of plants and the advanced Hi-C methods have rarely been reported in plants. Due to proximity ligation strategy in Hi-C, noise could be generated during the process (Kong and Zhang, 2019). Extracting reliable gene loops from the high-noise Hi-C data usually requires deep sequencing and significant bioinformatics efforts (Kong et al., 2020). Hi-C often needs substantial starting material (3∼5 g leaf) to obtain enough sequencing depth, making constructing a library infeasible for multiple samples (Yadav et al., 2021a). Thus, the high noise of Hi-C technology impede the development of 3D plant genomics.

In this study, we showed BL-Hi-C excellent performance in genome-wide profiling of chromatin interactions in *B. rapa* and *B. oleracea*. Compared with traditional Hi-C, BL-Hi-C requires significantly less sample input and lower costs. Moreover, we observed that BL-Hi-C exhibits decreased background noise and increased signal, which enhances its reliability and accuracy in detecting gene loops. By applying the BL-Hi-C to *B. rapa*, we delineated the genomic architecture and observed the configuration of *B. rapa* chromosomes. Our findings highlight the potential of BL-Hi-C as a valuable tool for investigating 3D genome organization in plants.

## RESULTS

### Modification of BL-Hi-C for plant species

Original BL-Hi-C requires a minimum of 0.5 million cells for the identification of chromosome interactions. However, counting the number of nuclei under the microscope can be challenging due to the presence of impurities in the nuclei isolated from plants (Figure S1B). Instead, we developed a semi-quantitative approach to estimate the number of nuclei by using the appropriate amount of plant leaves. Our results showed that 1 g of fresh leaves of *B. rapa* was sufficient to obtain 0.5 million nuclei. Traditional Hi-C usually requires 3∼5 g of fresh leaf. This reduction in starting material not only simplifies the experimental procedure but also allows the experiment to be completed within 2.5 days in a 1.5 ml tube (Figure 1A). Additionally, by reducing the reaction volume, the cost of library generation was able to decrease to as low as $92 per sample, which is approximately one-third the cost of traditional Hi-C (Table 5).

**Figure 1.**
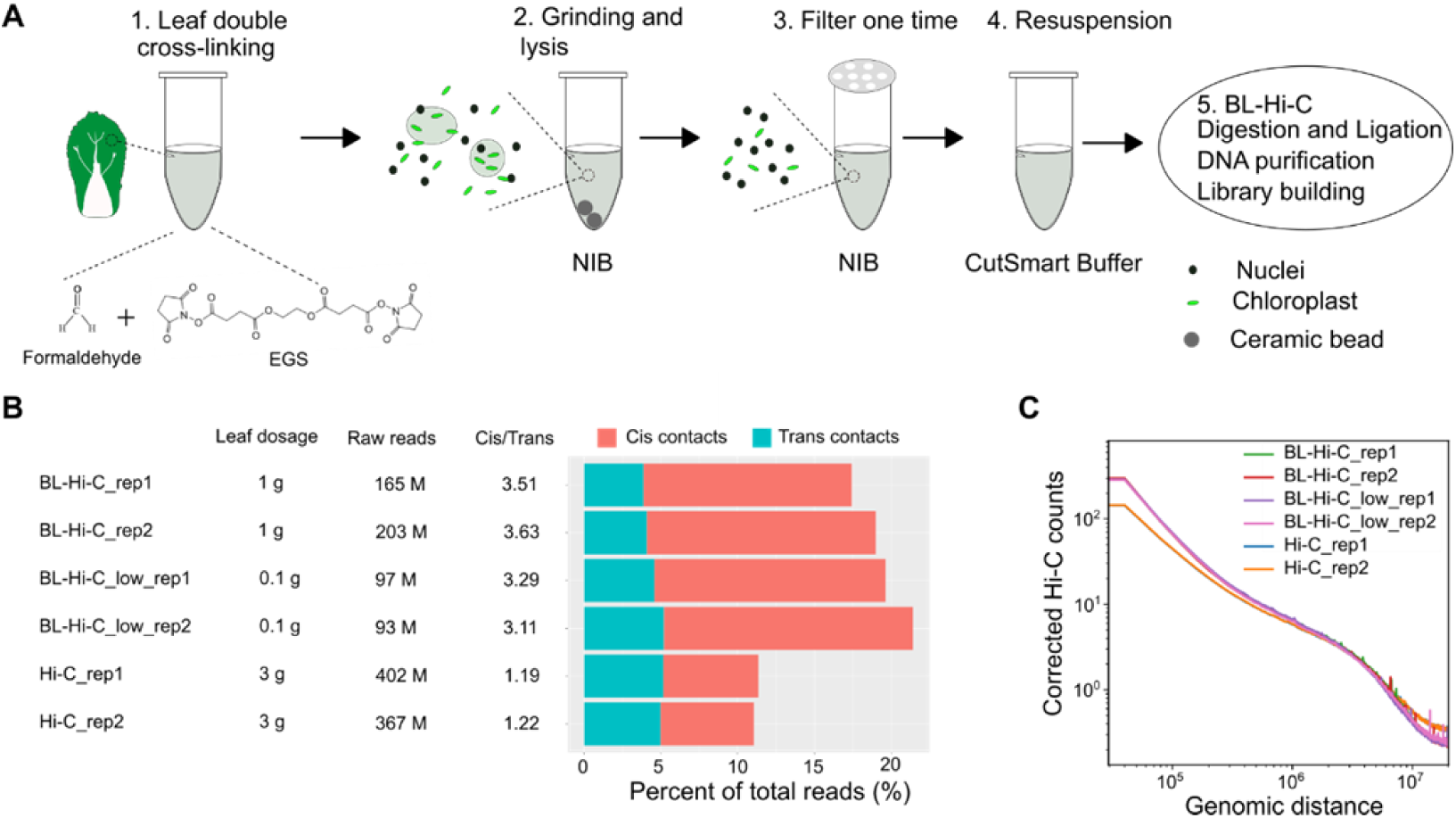
Schematic of BL-Hi-C method. (A) The nuclei extraction procedure includes double cross-linking, grinding and lysis, filter, and resuspension. (B) Efficiency comparison of BL-Hi-C, low input BL-Hi-C and Hi-C (Xie et al., 2019) in *B. rapa*. The cis contacts refer to intrachromosomal interactions, and the trans contacts refer to interchromosomal interactions. (C) Chromatin contact frequency (y-axis) was plotted as a function of linear genomic distance (x-axis) measured by BL-Hi-C and Hi-C in *B. rapa*.

We simplified the nuclei extraction procedure. In BL-Hi-C, the digestion and ligation processes are completed within the intact nuclei. Previous nuclei isolation protocols require a significant amount of tissue and long duration for completion because of grinding samples manually under liquid nitrogen and the multiple purification steps. We simplified the nuclei extraction procedure by grinding the leaf tissue with an electric grinder (step 2, Figure 1A) and reducing the filtration step to only once (step 3, Figure 1A). After resuspension in CutSmart Buffer (step 4, Figure 1A), nuclei were used to construct the library following the standard BL-Hi-C procedure (step 5, Figure 1A, Figure S1A).

To ensure that all steps of the BL-Hi-C library construction had been performed correctly, DNA fragments from each step were purified and analyzed by gel electrophoresis (Figure S1C). A typical smear of DNA fragments was observed after chromatin digestion using the restriction enzyme *Hae* III. After ligating digested chromatin to linkers, the DNA fragments were aggregated. These results indicate that the genome digestion and the proximity ligation were completed.

### Consistent key feature of genome organization captured by BL-Hi-C and Hi-C

We performed BL-Hi-C using the optimized protocol for *B. rapa*, generating 165 and 203 million sequencing reads in two *B. rapa* biological replicates. To further assess the power of BL-Hi-C in 3D genome structure analysis for plant species, we compared the 3D genomic data of *B. rapa* generated by Hi-C (Xie et al., 2019) and BL-Hi-C using the same data analysis pipeline. After mapping and data filtration, 28.8 and 38.7 valid contacts (non-redundant reads with PE ends mapped to different digested fragments, Supplementary Table 1) were obtained for each BL-Hi-C replicates. The ratio of valid contacts with respect to sequencing reads is 18%, which is higher than that of Hi-C (11%) (Figure 1B). The ratio of uniquely mapped reads of BL-Hi-C (∼30%) was significantly higher than that of Hi-C (∼12%) after filtering low-quality reads (MAPQ < 10). This result suggests that bridge linkers can help split chimeric fragments, thereby improving the ratio of uniquely mapped reads. In addition, BL-Hi-C possessed cis:trans ratio as high as 3.5 compared with 1.2 in Hi-C (Figure 1B), which indicated the BL-Hi-C library quality was substantially improved. Next, we employed HiCExplorer (Wolff et al., 2018) to quantitatively assess BL-Hi-C and Hi-C libraries. We found that BL-Hi-C libraries were highly reproducible among two methods (Figure S2, Pearson correlation coefficient = 0.9). Besides, BL-Hi-C identified chromatin contacts over a wide range of distances with an efficiency comparable to that of the Hi-C method (Figure 1C). To assess the robustness of BL-Hi-C, we conducted the assay using leaves of *B. oleracea*. A total of 314 million sequencing reads was generated in *B. oleraceaa* and 64.6 million valid contacts were obtained occupying 20% of total reads (Table S1). The ratio of cis:trans is 3.09 in BL-Hi-C which is higher than that in Hi-C (1.13) (Xie et al., 2019). The results of the experiment once again demonstrated the excellent performance of BL-Hi-C.

In order to validate the ability of BL-Hi-C in detecting crucial aspects of genome architecture, we conducted a comparative analysis of the contact heat maps generated by the two methods (Figure 2A, Figure S3). Despite being derived from distinct sequencing depths in *B. rapa*, the heat maps exhibited analogous interaction patterns at multiple scales, including the genome, chromosome, and local levels. Moreover, both methods consistently revealed a higher prevalence of cis-interactions compared to trans-interactions, as well as a higher frequency of interactions within chromosome arms compared to inter-arm interactions. In *B. oleracea*, BL-Hi-C and Hi-C also showed similar interaction signals across the whole genome (Figure S5).

**Figure 2.**
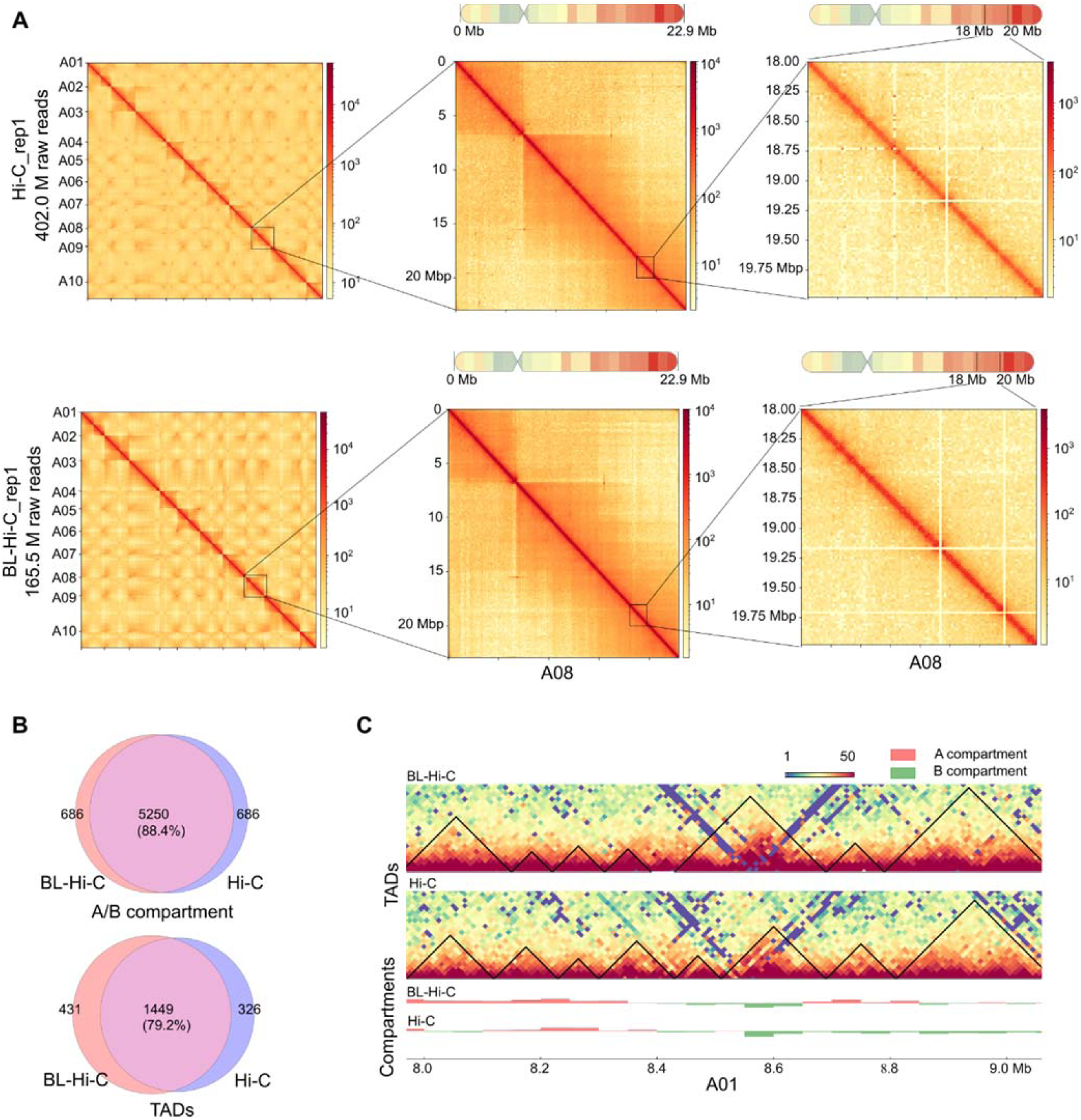
Results comparison between BL-Hi-C and Hi-C in *B. rapa*. (A) Heatmaps reconstructed using data generated from Hi-C, BL-Hi-C and low-input Hi-C. The resolution was set as 500 Kb for the entire genome, 150 Kb for the A08 chromosome, and 20 Kb for the local regions. (B) Overlap of compartments and TADs between BL-Hi-C and Hi-C. (C) Part of genome browser images showing TAD structure detected on A01 by BL-Hi-C and Hi-C.

In 3D genomics, compartments and TADs play critical roles in regulating gene expression. We compared the compartments and TADs obtained from BL-Hi-C and Hi-C employing the same analysis pipeline (Table 2, Table 3). The BL-Hi-C and Hi-C displayed a high degree of consistency regarding their A/B compartments. About 88.4% of bins shared the same compartment status (Figure 2B). Here only TADs from BL-Hi-C and Hi-C overlapped with each other more than 80% were considered overlapped TADs. We identified 1,880 TADs from 368 million raw reads in the BL-Hi-C, 1,449 (77.1%) of which were shared with TADs in Hi-C. Besides, the visualization result of TADs showed TADs in BL-Hi-C were very similar to those in Hi-C (Figure 2C). Taken together, BL-Hi-C faithfully captures the key features of genome architecture in the plants evaluated.

### Low background noise of BL-Hi-C

To assess the signal-to-noise of BL-Hi-C relative to Hi-C, we compared interaction profiling generated by BL-Hi-C and Hi-C in *B. rapa* (Figure 3A). We observed that the BL-Hi-C signals were much clearer and sharper than Hi-C by genome browser visualization. Therefore, the detected chromatin interactions from BL-Hi-C were more distinct than that of Hi-C. We next identified 140,944 overlapped peaks between biological replicates in BL-Hi-C (Figure 3B, Table 4). The percentage of overlapped peaks was higher in BL-Hi-C (97.92%) than that in Hi-C (88.87%), which identified 131,405 overlapped peaks between biological replicates (Figure 3B). The result highlights the better peak reproducibility of BL-Hi-C. A total of 58,968 common peaks can be found between BL-Hi-C and Hi-C (Figure S4). On average, there were average 2.3 normalized interactions per BL-Hi-C peak, which was significantly higher than 1.4 identified from Hi-C data (p-value < 2.2e-16, Figure 3D). BL-Hi-C peaks also had a higher-log(qvalue) than Hi-C peaks (p-value < 2.2e-16, Figure 3E). To further evaluate the signal, we compared the average read counts for BL-Hi-C and Hi-C datasets around the peaks. We found that BL-Hi-C profiling showed substantially more signal accumulation at peak summits, implying that BL-Hi-C would be more effective in distinguishing interaction (Figure 3F). The result implied that BL-Hi-C peaks were separated from each other. By visual examination, we also found that interactions detected by BL-Hi-C were more concentrated around the peaks (Figure 3A). Similar interaction signals were also observed in *B. oleracea* (Figure S6A). BL-Hi-C also showed higher reads density in peaks and higher signal accumulation at peak summits compared with Hi-C in *B. oleracea* (Figure S6B, S6C). These results demonstrated that BL-Hi-C had the advantage over Hi-C in signal-to-noise ratio for chromosome conformation capture.

**Figure 3.**
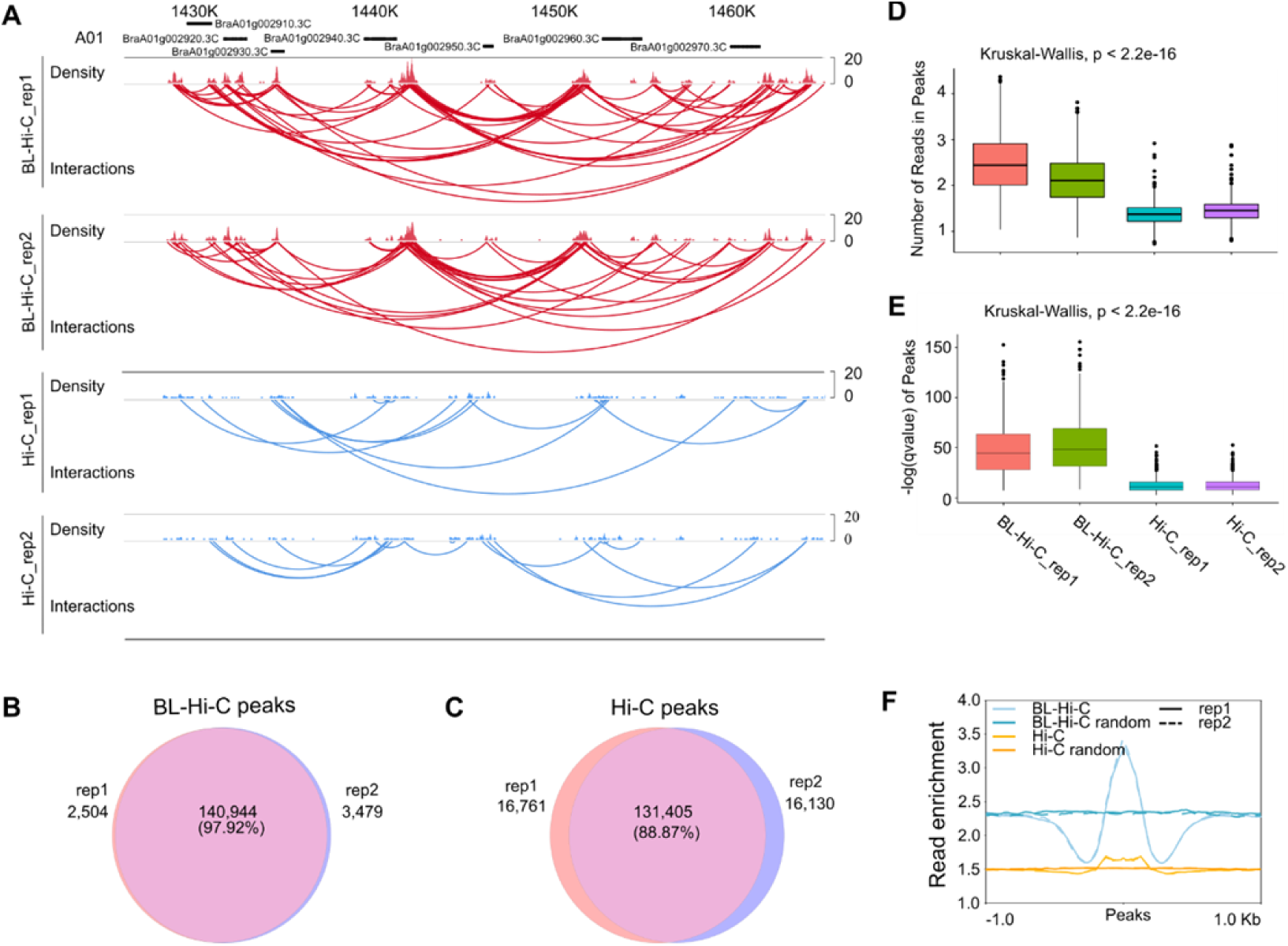
Distribution of interaction signals around peaks detected by BL-Hi-C and Hi-C in *B. rapa*. (A) Interaction signals in a local region of chromosome A01. (B-C) Venn diagram showing the overlapped peaks of two biological replicates. The percentage was calculated by common peaks divided by average peaks. (D-E) Boxplots showing the number of reads falling on peaks and-log(qvalue) of peaks. (n=1000). (F) Metaplot of signal enrichment around peaks. The randomly selected regions were taken as background.

We evaluated the sensitivity of BL-Hi-C by performing a comparative analysis of the gene loops identified at the *FLC* loci between BL-Hi-C and Hi-C. Previous studies have reported the presence of a gene loop between the *FLC* promoter and the downstream in *Arabidopsis* (Crevillen et al., 2013; Liu et al., 2016). In *B. rapa*, there are four *FLC* homologs: *BrFLC1*, *BrFLC2*, *BrFLC3*, and *BrFLC5* (Schranz et al., 2002). Our BL-Hi-C analysis of Chiifu revealed obvious gene loops in *BrFLC1*, *BrFLC2*, and *BrFLC3,* including loop types of promoter-downstream, promoter-intron and intron-downstream (Figure 5). However, such gene loops were not observed in the Hi-C analysis. Neither BL-Hi-C nor Hi-C detected gene loop within the *BrFLC5* region, indicating that *BrFLC5* may not form gene loop. This is consistent to the fact that the *BrFLC5* allele in Chiifu is non-functional (Xi et al., 2018). Similarly, in JZS (*B. oleracea*), the BL-Hi-C analysis detected obvious gene loops within *BoFLC1*, *BoFLC2*, and *BoFLC3* (Figure S6D). These findings highlight the high sensitivity of BL-Hi-C in detecting gene loops, largely owing to its low background noise.

### Low input of BL-Hi-C

The Hi-C technique is limited in its applicability for rare cell research due to its requirement for a substantial amount of leaf material, which also complicates the experimental process and hinders high-throughput library building. The improved performance of BL-Hi-C suggested that this method might work efficiently with limited samples. To test the feasibility of a small amount of sample using BL-Hi-C, we built the BL-Hi-C library using only 100 mg of leaf. With this low input library, we were able to generate 93 and 97 million raw read pairs in two biological replicates, respectively. Notably, the proportion of valid contacts (21.1%) and the ratio of cis to trans contacts (3.3) in low-input BL-Hi-C were similar to those of normal input BL-Hi-C (Figure 1B). Additionally, low-input BL-Hi-C showed similar interaction patterns with BL-Hi-C in heatmaps (Figure S3).

To characterize the peaks identified through low-input BL-Hi-C, we conducted a comparative analysis with BL-Hi-C data. Visualizations of the genome browser showed high consistency between the BL-Hi-C and low-input BL-Hi-C contact signals (Figure 4A). We further identified 121,303 and 123,155 peaks in two biological replicates of low input BL-Hi-C, respectively. Of these, 117,095 peaks were commonly observed, accounting for 95.8% of the total (Figure 4B). Notably, 99% of the low-input BL-Hi-C peaks overlapped with the BL-Hi-C peaks (Figure 4C). It was evident that low input library reads enriched around peaks, which was similar to BL-Hi-C (Figure 4B). The high reproducibility (Spearman’s correlation= 0.986) of the low-input BL-Hi-C peak intensity between the two replicates was demonstrated (Figure 4D). Additionally, a high correlation (Spearman’s correlation=0.963) was also observed between low-input BL-Hi-C and BL-Hi-C in peak intensity (Figure 4D), confirming that decreasing the input amount to as low as 100 mg tissue did not reduce the roubustness of BL-Hi-C.

**Figure 4.**
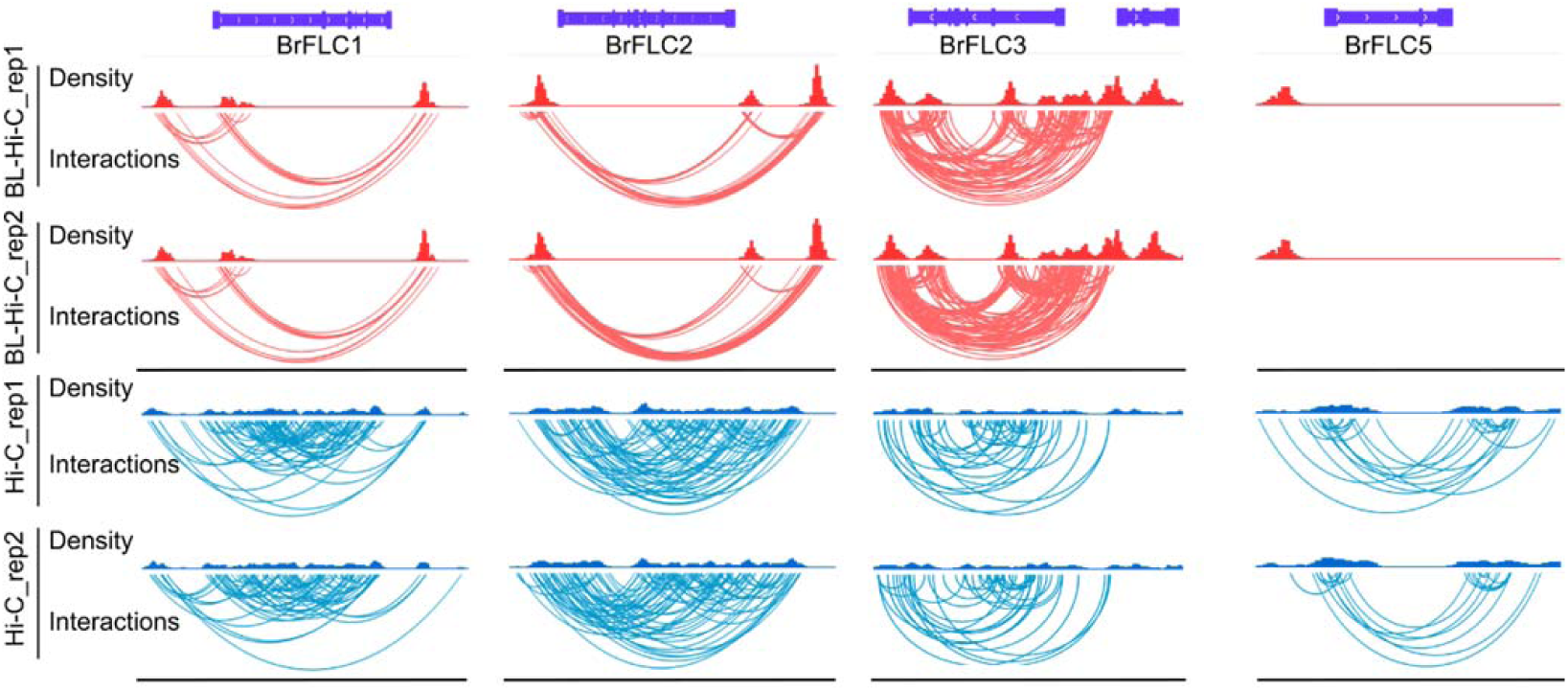
Gene loops at BrFLC1 (BraA10g027720.3C), BrFLC2 (BraA02g003340.3C), BrFLC3 (BraA03g004170.3C) and BrFLC5 (BraA03g015950.3C) revealed by BL-Hi-C and Hi-C.

**Figure 5.**
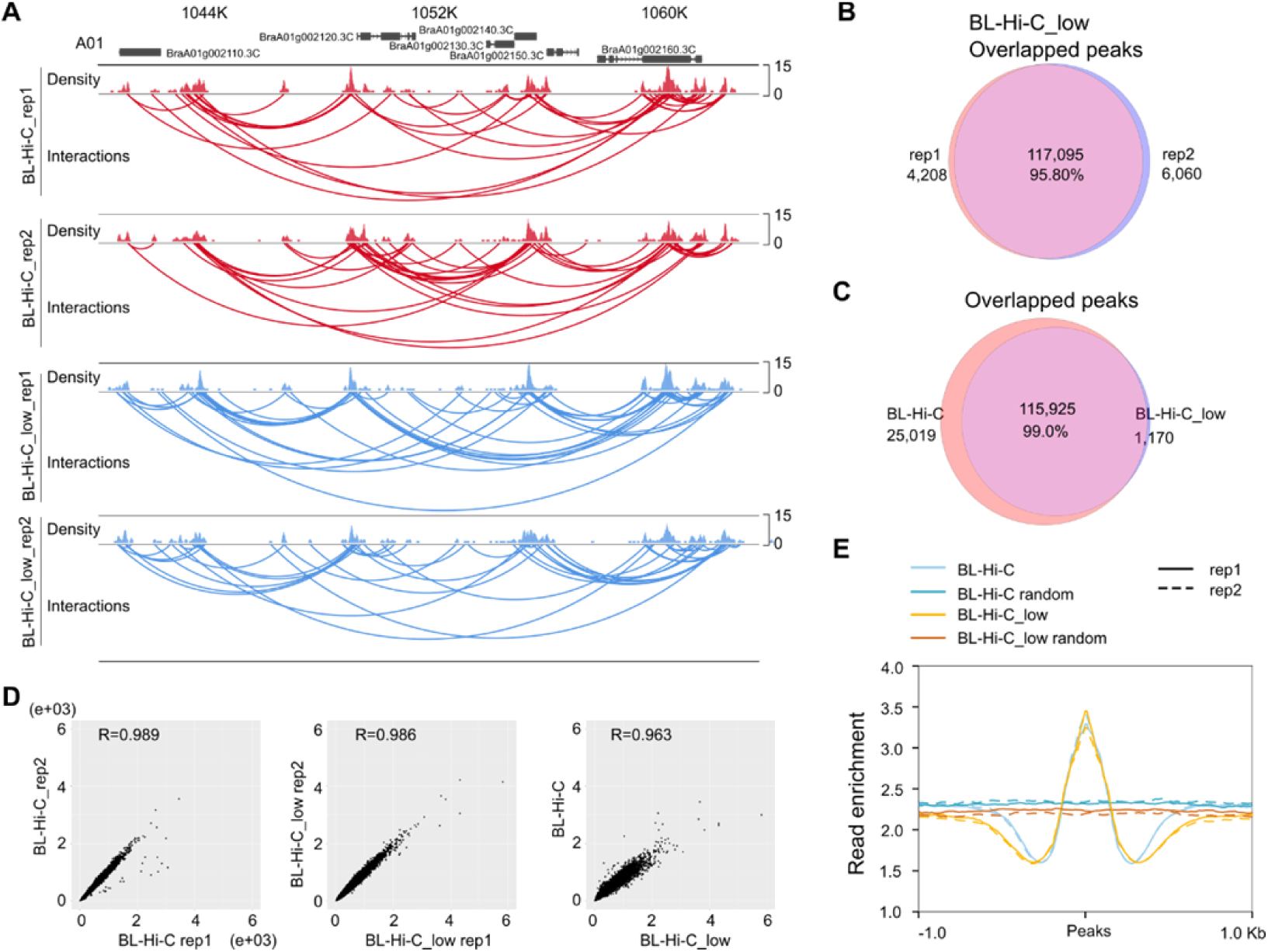
Comparision of peaks detected by BL-Hi-C and low input BL-Hi-C in *B. rapa*. (A) Interaction signals in a local region of chromosome A01. (B-C) Venn diagram showing the overlapped peaks of biological replicates and two methods. The percentage was calculated by common peaks divided by average peaks. (D) Scatter plots of the peak intensity between different datasets’ peak (n=144,832) loci. The R-value is the Spearman’s correlation coefficient. (E) Metaplot of signal enrichment around peaks. The randomly selected regions were taken as background.

**Figure 6.**
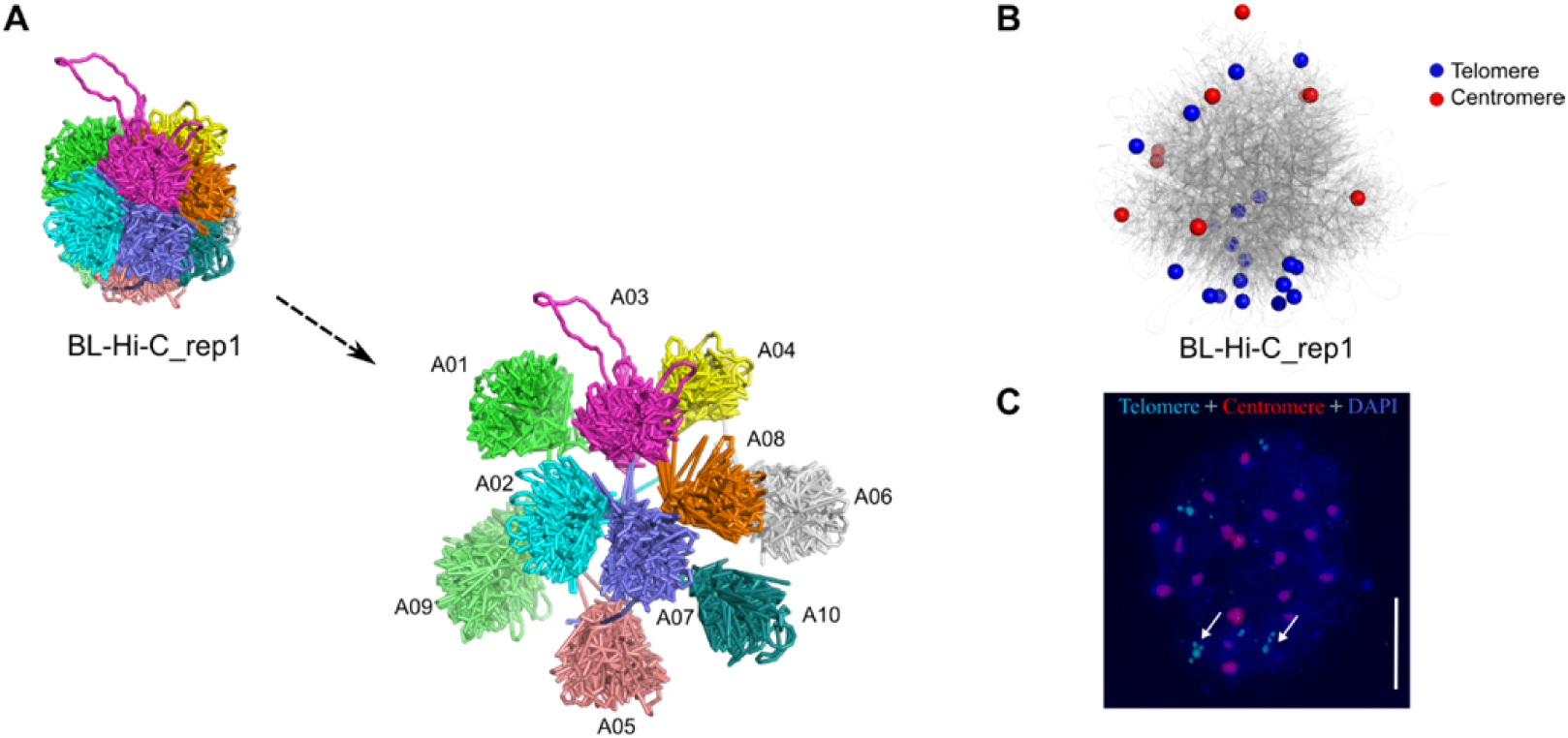
Reconstructed particle-on-a-string 3D genomes of *B. rapa*. (A) 3D organization with expanded views of the separate chromosome territories. (B) Spatial distribution of telomeres and centromeres of ten chromosomes in the 3D genome. The blue sphere indicates the telomeres, and the red sphere indicates centromeres. Each particle equals ten kilobase pairs. (C) Chromatin was stained blue with 4 ′,6-diamidino-2-phenylindole (DAPI). Fluorescence in situ hybridization was performed with probes specific for centromeres (red) and telomeres (blue). Arrows indicate clusters of multiple telomeres. Scale bar, 2 μm.

### Delineating the Bouquet configuration of *B. rapa*

To investigate the chromosome organization in *B. rapa* nucleus, we constructed 3D genome structures (Figure 5A) using a particle-on-a-string representation and the extended simulated annealing software Nuc_dynamic (Stevens et al., 2017). Our analysis of the 3D genome architecture of *B. rapa* nuclei illustrated that each chromosome occupied an exclusive region within the nucleus, supporting the concept of “chromosome territory”.

We observed Bouquet configuration in *B. rapa* cells. Intrigued by the morphology of *B. rapa* nuclei, we labeled the ten centromeres and twenty telomeres on the 3D structural model of the genome with different colors. The telomeres were found to be localized close to each other, while the centromeres were located at the periphery of the nucleus (Figure 5B). We observed similar results in *B. oleracea* as well (Figure S7). Notably, one of the A03 telomeres locates on the surface rather than inside the 3D topological structure (Figure S8A). Then, we observed the position of the centromeres through different angles of the 3D genome model (Figure S8B), which revealed that the centromeric region was visible on the side of the nucleus. Fluorescence *in situ* hybridization also supported clusters of telomeres (Figure 5C). These results demonstrate the presence of the Bouquet configuration in *B. rapa* leaf nuclei.

## DISCUSSION

Over the past decade, traditional Hi-C has significantly advanced research on 3D genomes (Pei et al., 2021). BL-Hi-C uses linker ligation to overcome the disadvantages of the traditional Hi-C such as inefficiency and insensitivity (Liang et al., 2017). In the present study, we simplified the nuclei extraction step to enable efficiently utilizing BL-Hi-C in plant research. Our BL-Hi-C result domonstrated that the simplified nuclei extraction method was robust (Figure S1B). It has the potential to be used in other technologies, such as plant ATAC-seq and CUT&Tag. Importantly, we found BL-Hi-C is simple, low noise, and low cost compared to traditional Hi-C, especially when construct library using low input material. Therefore, BL-Hi-C has great potential in plant 3D genomics research.

The analysis of a few samples may not be suitable for understanding the regulatory principles of chromatin structure due to chromatin interaction is very dynamic (Yadav et al., 2021a). Researchers had been limited to analyzing only a few samples using Hi-C due to complex library construction procedures and high cost. A fast and low-cost library construction procedure is necessary for analysis of chromatin interactions in a large number of samples. Though commercial kits for Hi-C provide better performance over traditional Hi-C. The significant expenses associated with commercial Hi-C kits also pose a substantial barrier to their widespread adoption in 3D plant genomic studies. Hi-C-based fluorescence-activated cell sorting (FACS) with a scale of approximately 1×10^5 nuclei has been employed to investigate plant 3D genomes (Yadav et al., 2021b). Due to technical limitations, current plant pan-3D genomes only enable the analysis of a small number of samples (less than 30).

Without purchasing costly instruments and performing complex procedures, we present that BL-Hi-C is a simple, cheap, and low-input feasible method that allows us to examine three-dimensional genomes in a high-throughput way. Low-input BL-Hi-C can also be effectively applied to analyze samples with limited starting material, such as pollon and shoot tips, which have received relatively little attention in research endeavors.

Previous investigations into the 3D genomes of plants have predominantly concentrated on compartments and topologically associating domains (TADs), with less emphasis on chromatin loops. In mammalian organisms, chromatin loops are believed to have a pivotal role in facilitating specific interactions and communication between enhancers and promoters (Oudelaar and Higgs, 2021). Nonetheless, the identification of chromatin loops in plants using Hi-C technology poses challenges due to high background noise, unless ultra-deep Hi-C datasets containing billions of contact reads are employed (Pei et al., 2022). The application of ChIA-PET technology, characterized by reduced background noise, proves to be advantageous for investigating chromatin loops in plants even at lower sequencing depths (Peng et al., 2019; Zhao et al., 2019). However, it should be noted that ChIA-PET analysis is limited to specific protein-mediated chromatin loops (Tang et al., 2015). An alternative approach, known as BL-Hi-C, employs an independent antibody enrichment method and exhibits lower noise levels compared to conventional Hi-C. Furthermore, our research demonstrates that BL-Hi-C can successfully detect gene loops, including promoter-downstream loops, promoter-intron loops, and intron-downstream loops, which were undetectable using Hi-C. Consequently, the utilization of BL-Hi-C has the potential to facilitate the exploration of gene loops in plants.

The spatial arrangement and functional properties of individual chromatin domains in nuclei play a critical role in 3D genomics. Traditional methods such as FISH and GFISH (Fransz et al., 2002) were previously used to explore spatial chromosome distribution. Although Hi-C has emerged as a powerful technology for revealing chromatin conformation, the results usually were based on heatmap observations, which is an indirect method (Mascher et al., 2017; Concia et al., 2020). In this study, we constructed a 3D genome model representing the Bouquet configuration and directly located the telomeres of *B. rapa* chromosomes in the middle and centromeres on the surface. Our method could also be used to visualize other 3D structures such as TADs and chromatin loops.

In summary, we showed BL-Hi-C technology exihibits stronger interaction signals compared to that of Hi-C, and the feasibility of low-input library construction. These results indicates BL-Hi-C is a powerful and widely applicable method for analyzing plant 3D genomes.

## MATERIALS AND METHODS

### Plant material

1. *B. rapa* (ssp. pekinensis, cultivar Chiifu) and *B. oleracea* (cultivar JZS) seeds were cultivated in Petri dishes at a home temperature (25 °C) for 14 h, and the seedlings were transferred to soil in a greenhouse at 23 °C with a 16 h photoperiod. After 1 month, the harvestable young leaves were used for BL-Hi-C.

### Experimental protocol for plant BL-Hi-C

### Double cross-linking

For *B. rapa* and *B. oleracea*, 2 g of material of each sample was collected in the 50 ml tube, adding 20 mL of NIB (20 mM Hepes (pH8), 250 mM sucrose, 1mM MgCl_2_, 0.5 mM KCl, 40% glycerol, 0.25% Triton X-100, 0.1 mM phenylmethanesulfonylfluoride (PMSF), 0.1% 2-mercaptoethanol), 20 ml of 4% formaldehyde (Sigma, #F8775) and 100 μl 0.15 M of EGS (Thermo, #21565) to submerged leaves. Then, the tube was placed in a desiccator, applying vacuum for 1 h. After cross-linking, the remaining formaldehyde was sequestered by adding 2680 μl of 2 M glycine (Sigma, G7126) and continued applying the vacuum for 5 minutes. Next, the NIB/formaldehyde mixture was removed, washing the leaf using ddH2O.

### Nuclei extraction

Samples were ground in liquid nitrogen to a fine powder and then lysed in NIB. The mixture was aliquotted into two tubes and filtered once with Miracloth (Millipore, #475855). After spinning down the filtrate at 3000 g at 4℃ for 15 mins, the supernatant was removed. Subsequently, the nuclei were resuspended in 1.3×CutSmart Buffer and spun down at 1900 g at 4℃ for 5 min.

### Nuclei lysis and restriction digestion

Nuclei were then resuspended in 50 µl of 0.5% SDS and incubated at 62°C for 10 mins on the thermomixer, shaking at 900 r.p.m. After nuclei lysis, 145 µl of ddH_2_O and 25 µl of 10% (v/v) Triton X-100 were added into the tube to quench the SDS reaction. The mixture was then gently shaken for 15 minutes at 37℃. Then, 25 µl of 10×CutSmart Buffer and 10 µl of *Hae III* (NEB, #R0108L) were added to each tube, and the tubes were incubated at 37°C for 12 h with rotation at 900 r.p.m. Finally, 2.5 µl of 100 mM dATP solution and 2.5 µl of Klenow Fragment (3′->5′ exo-) (NEB, #M0212L) were added to the mixture, and the mixture was incubated for 40 minutes at 37℃ with rotation at 900 r.p.m.

### Proximity ligation

After restriction enzyme digestion, the linker (F:pCGCGATATC/iBiodT/TATCT GACT, R:pGTCAGATAAGATATCGCGT) was ligated to the digested chromatin. In each tube, 750 µl of ddH_2_O, 120 µl of 10× T4 DNA ligase buffer, 100 µl of 10% (v/v) Triton X-100, 5 µl of T4 DNA ligase (NEB, #M0202L), and 4 µl of 200 ng/µl linker were added to the 260 μ l of digested chromatin and mixed thoroughly. The mixture was then incubated at 16 °C for 4 h with rotation at 900 r.p.m. After linker ligation, chromatin DNA-protein complexes were centrifuged at 3500 g for 5 minutes at 4°C, and the supernatant was discarded. Next, the pellets were resuspended in 309 µl of ddH_2_O, 35 µl of Lambda Exonuclease Buffer, 3 µl of Lambda Exonuclease (NEB, #M0262L), and 3 µl of Exonuclease I (NEB, #M0293L). The mixture was then incubated at 37°C for 1 hour at 900 r.p.m.

### DNA Purification

After proximal ligation, 45 µl of 10% SDS and 55 µl of 10 mg/ml proteinase K (Solarbio, #P1120) was added to the tube, incubated the nuclei at 60℃ for about 3 hours to reverse crosslinking. After incubation, 450 µl of phenol:chloroform:isoamyl alcohol (25:24:1) was added to the tube, which was shaken vigorously and then centrifuged for 15 min at 14,000 r.p.m. Next, 400 µl of supernatant was transferred into a new tube. DNA was precipitated with 400 µl of isopropanol, 40 µl of 3 M sodium acetate (pH 5.2), and 4 µl of Dr. GenTLE Precipitation Carrier (Takara, #9094) and centrifuged for 15 min at 12000 r.p.m. The precipitated DNA was washed once with 80% ethanol and dissolved in 40 μl of 0.1×TE Buffer.

### Library generation

The Bioruptor (Diagenode) was used to break the DNA into 300-500 bp using the following settings: Duty cycle 32; 30s on, 30s off. And then 1.2×Ampure XP beads (Beckman, #A63881) were used to purify the DNA fragments. After the pull-down of biotin-labeled DNA reads (Thermo, #11205D), the DNA Library Prep kit (Enzyme, ND608) for Illumina was used to complete DNA damage repair, end-repair, adaptor ligation, and PCR library amplification. After 12 amplification cycles, the DNA product was purified by 1×Ampure XP beads for deep sequencing.

### Analysis of BL-Hi-C

The trimLinker of ChIA-PET2 (v0.9.3)(Li et al., 2017) was used to filter the linker, and HiC-Pro (v3.0.0)(Servant et al., 2015) was used to align the sequence to the reference genome (*Brassica rapa* v3.0;http://39.100.233.196:82/download_genome/rassica_Genome_data/Brara_Chiifu_V3.0/Brapa_sequence_v3.0.fasta.gz; *Brassicaoleraceaa* v2.0; http://39.100.233.196:82/download_genome/Brassica_Genome_data/Braol_JZS_V2.0/Brassica_oleraceaa_JZS_v2.fasta.gz). Reads with low mapping quality (MAPQ < 10) was filtered out and reads with the same coordinate on the genome or mapped to the same digestion fragment was removed. The ICE method was applied to normalize the interaction matrix for different resolutions (10 Kb, 20 Kb, 40 Kb, 150 Kb, and 500 Kb). HiCExplorer (v2.1.4)(Wolff et al., 2018) was used to convert normalized matrix data into h5 format and other formats for further analysis. The Pearson correlation coefficients and interaction matrix were visualized using HiCExplorer. We combined biological replicate data for analysis of compartments and TADs. A/B compartments were determined by Juicer (v1.9.9) (Durand et al., 2016) at 50 Kb resolution. The TAD boundaries were analyzed by HiTAD at 10 Kb resolution (v0.4.2)(Wang et al., 2017). Peaks were called from valid contacts by MACS2 (v2.2.7.1) with default parameters. After normalizing with CPM, read enrichment was visualized by the WashU Epigenome Browser (Li et al., 2019). According to the previous research in rice single-cell (Zhou et al., 2019), 10 kilobase pixels were chosen as parameters for Nuc_dynamic (https://github.com/tjs23/nuc_dynamics) analysis. 500,000 valid contacts were randomly selected to construct the 3D model three times. Simulated annealing was calculated by the Nuc_dynamic software (parameter: −s 8 2 1 0.5 0.2 0.1 0.05 0.02 0.01) to create a PDB file for viewing the 3D genome structures in pymol (v4.60).

### Experimental protocol for low-input plant BL-Hi-C

### Double cross-linking

For *B. rapa*, 0.1 g of material of each sample was collected in the 1.5 ml tube, adding 500 μl of nuclei isolation buffer (NIB: 20 mM Hepes (pH8), 250 mM sucrose, 1mM MgCl_2_, 0.5 mM KCl, 40% glycerol, 0.25% Triton X-100, 0.1 mM phenylmethanesulfonylfluoride (PMSF), 0.1% 2-mercaptoethanol), 500 μl of 4% formaldehyde (Sigma, #F8775) and 10 μl 0.15 M of EGS (Thermo. #21565) to submerged leaves. Then, the tube was placed in a desiccator, applying vacuum for 1 h. After cross-linking, the remaining formaldehyde was sequestered by adding 67 μl of 2 M glycine (Sigma, G7126) and continued applying the vacuum for 5 minutes. Next, the NIB/formaldehyde mixture was removed, washing the leaf using ddH2O.

### Nuclei extraction

The electric grinder was used to grind the leaf with the parameter 60 s, 60 Hz, 4 cycles, and then lysed in NIB. The mixture was transferred into new tubes and filtered once with Miracloth (Millipore, #475855). After spinning down the filtrate at 3000 g at 4℃ for 15 mins, the supernatant was removed. Subsequently, the nuclei were resuspended in 1.3×CutSmart Buffer and spun down at 1900 g at 4℃ for 5 min. The next procedure was the same as that described above for the plant BL-Hi-C protocol.

## Data availability

All sequencing data generated for this study have been submitted to the NCBI Sequence Read Archive under accession number PRJNA945226. Previously published Hi-C data analyzed in this study can be obtained from GEO via accession code (SRR8633037, SRR8633038).

## Authors’ contributions

X.W. and J.W. designed the project; L.P.Z., R.Z. and L.Z. prepared materials and performed the experiments; L.P.Z., H.G. and Z.Z. performed the data analysis; L.P.Z., X.W. and J.W. wrote the manuscript; J.W., J.L. and X.C. revised the manuscript. All authors read and approved the final manuscript.

